# Methodological considerations for behavioral studies relying on response time outcomes through online crowdsourcing platforms

**DOI:** 10.1101/2023.06.12.544611

**Authors:** Patrick A. McConnell, Christian Finetto, Kirstin-Friederike Heise

## Abstract

This study explored challenges associated with online crowdsourced data collection, particularly focusing on longitudinal tasks with time-sensitive outcomes like response latencies. The research identified two significant sources of bias: technical shortcomings such as low, variable frame rates, and human factors, contributing to high attrition rates. The study also explored potential solutions to these problems, such as enforcing hardware acceleration and defining study-specific frame rate thresholds, as well as pre-screening participants and monitoring hardware performance and task engagement over each experimental session. This study provides valuable insights into improving the quality and reliability of data collected via online crowdsourced platforms and emphasizes the need for researchers to be cognizant of potential pitfalls in online research.

## Introduction

Relying on human intelligence is essential to infer neural mechanisms underlying human behavior. This insight has not changed since the invention of the original Mechanical Turk ^1^ and has regained recent attention, for instance, in the field of neuroscience ^2,3^. As the Mechanical Turk relied on human chess masters to operate it, so does human behavioral research depend on human participants. Over the past decade, experimental behavioral research has increasingly turned to online crowdsourcing to recruit study participants, as evidenced by a steep increase in PubMed-listed publications. With closed-down labs, this trend accelerated since the start of the COVID-19 pandemic. The number of publications mentioning “Amazon Mechanical Turk” and “MTurk”, one of the most popular platforms for crowdsourced research, increased from < 10 per year in 2013 to ∼ 200 per year in 2022, doubled from pre-to post-pandemic, and continues to increase. Online experimental designs accessible through MTurk range from simple surveys to labor-intensive experimental designs like probabilistic learning ^4^ and cognitive control task batteries ^5^. The versatility of online research tools using platforms like MTurk additionally highlights this trend and its appeal to researchers across many fields ^6^.

Among many experimental paradigms in psychophysics, studying response times constitutes the primary outcome to infer underlying cognitive control processes and assess changes in cognitive functioning ^7^. The precise evaluation of response latencies is of particular interest, especially when investigating behavioral change over time, for example, to study learning and memory ^8–10^. Unfortunately, several significant methodological shortcomings have been reported ^11–14^, which cause considerable obstacles to interpreting reaction time measurements collected online. Additionally, significant time-of-day variations in demographic composition ^15^, inadequate worker comprehension of task instructions ^16^, worker inattentiveness ^17,18^, non-random attrition (i.e., loss of participants over time) ^19^, bots and poor data quality ^20^ have been documented. We here add to this list of potential pitfalls with a focus on the reliability of timing-based outcome parameters.

Based on preliminary findings from an ongoing longitudinal procedural learning study using MTurk, we expand upon the known limitations of online crowdsourcing data and specifically shine a light on the challenges resulting from longitudinal data collection relying on time-sensitive outcomes. Navigating circumstances such as these is of high relevance to research involving psychophysics broadly, and therefore, we discuss two major points, which we consider essential in this context: technical difficulties arising from hardware and software configurations and complications arising from human factors. In discussing the implications of these hurdles, we want to motivate a discussion about potential solutions for improving data quality and provide guidance for researchers planning to use online crowdsourcing platforms to conduct psychophysics research.

## Methods and Results

### Overview of the Experimental Design and Data Collection Process

Response times were collected through an experiment that was created using PsychoJS and then hosted on Pavlovia (https://pavlovia.org/). Participants were recruited via “Human Intelligence Tasks” [HITs] on MTurk (https://www.mturk.com/) and given access to the experiment on Pavlovia. Access to MTurk HITs was restricted to workers with > 95% prior HIT approval ratings with and located in countries within Europe and North America at the time of screening.

Briefly, participants (i.e., workers) went through seven experimental sessions preceded by a screening session (session 0) in which the experiment was described and informed consent was actively obtained according to approved local Institutional Review Board protocol. Additionally, the screening session was used to filter out bots and inattentive workers based on a simple question testing an obvious detail of the described study procedures. The experimental paradigm required participants to match visual cues to keys on their keyboard and prompted them to respond with their best possible temporal precision. Behavior was tracked through response times of key presses and releases indexing performance precision, which was used to compute behavioral change over time across consecutive sessions.

Participants were informed about requirements for admittance to subsequent sessions and reimbursement upon compliance, defined as adhering to the instructions, and consistency of performance. Non-compliance was explicitly defined as reason for rejection whereas level of performance was specified as requirement for qualification (i.e., admittance) to the subsequent session. Workers received payment only for non-rejected sessions. Each of the seven experimental sessions, which were approximately 20-25 minutes in length, were published on MTurk around noon US Eastern Standard Time [EST]. All sessions were required to be completed within 24 hours of publishing in order to gain access to the respective next session. Performance data were manually reviewed daily for task compliance, and submissions were approved and qualified or rejected depending on criteria mentioned above.

### Point 1. Technical Factors Contributing to Attrition

Given their impact on response time precision, we closely monitored participants’ hardware frame rates throughout the study. While relying on PsychoJS via Pavlovia to calculate frame rates at the beginning of the experiment, we moved to a custom calculation method that involved averaging the frame times during each experimental trial. The advantage of this method was that it allowed the examination of frame rate variability within and across experimental sessions. We here report the custom frame rate calculations only; for completeness, comparison with the frame rate implemented through Pavlovia is presented in supplemental online information available at https://doi.org/10.6084/m9.figshare.23499099.

We realized a significant variation in frame rate precision with software version updates. Specifically, after updating to a newer version of PsychoJS (i.e., from 2021.2.3 to 2022.2.4), a significant number of sessions with very low frame rates (below 20 frames per second [fps]) were evident. To address this problem, we utilized a custom PsychoJS version (2022.2.4 custom) provided by Pavlovia that allowed detecting and logging graphics processing unit [GPU] based hardware acceleration. Only through this step, it was possible to identify cases in which GPU hardware acceleration was disabled, and to automatically switch to an alternative renderer optimized for central processing unit [CPU] rendering. As shown in our results reported below, enabling hardware acceleration is critical for acquiring reliable and accurate response times. In contrast, disabled hardware acceleration results yield untrustworthy data (i.e., low and variable frame rates), which in our case led to inacceptable data quality and hence discarded crowdsourced data.

Of a total of 4028 sessions across all 7 experimental days, 119 sessions were run with PsychoJS 2021.2.3, and 3909 were run with PsychoJS 2022.2.4. Hardware acceleration information was available for a subset of these sessions (n = 3781); only 499 (13.20%) were run with hardware acceleration and the vast majority, i.e., 3282 (86.80%), were run with hardware acceleration disabled. Hardware acceleration information was available for 781 unique workers. Of these, 683 workers (87.5%) did not have hardware acceleration enabled, while only 98 (12.5%) did. While the sample running with hardware acceleration also showed lower frame rates in some cases, it was overall considerably more consistent, with most of the data points at 60 fps and a small number at higher frame rates (144 fps) (Fig. 1).

**Figure 1.**
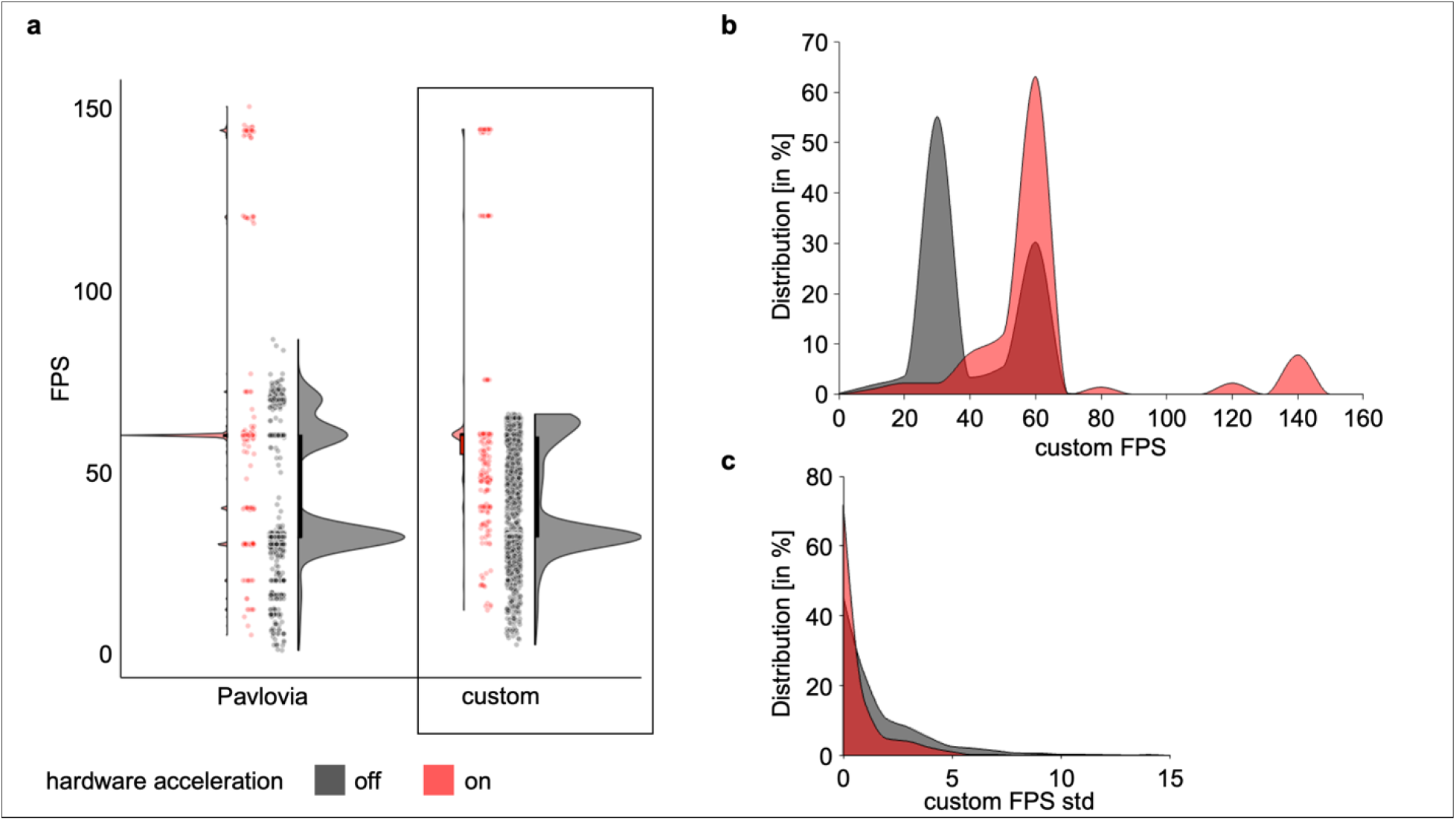
Hardware acceleration influences on frame rate and frame rate variability. Hardware acceleration disabled (‘off’) shown in black, enabled (‘on’) depicted in red. **a) Comparison of Pavlovia and custom frame rate calculations**. Frames per second (FPS) shown for both frame rate sources, i.e., Pavlovia’s built-in (left) that sampled one data point per session onset, and custom (right), which sampled frame rate across trials within session. **b) Normalized distributions of frame rate**. Depicts effects of hardware acceleration on custom frame rate calculation across all available data with hardware acceleration ‘on’ showing higher and less variable frame rates than ‘off’. **c) Variability distributions of custom frame rate**. Depicts within session, between-trial variability (standard deviation, std) of custom frame rate for individual workers with hardware acceleration ‘on’ showing less variability within session than ‘off’. Data and code to reproduce the figures are available at https://doi.org/10.6084/m9.figshare.23499195.

### Point 2. Human Factors Contributing to Attrition

We tracked the number of participants who passed through the various stages of screening and approval, as well as those who failed to submit data or were rejected based on task performance. Out of the 1820 participants who actively provided informed consent, 137 (7.53%) failed to pass the initial screening question qualifying for admission to the first experimental session. Thus, a total of 1683 (92.47%) workers were eligible to proceed to session 1. The highest drop-out occurred during session 1 due to non-compliance with task instructions, resulting in only 784 participants (43.08%) qualifying to move forward to session 2. Across sessions 2-7, further non-compliance (i.e., due to lost-to-follow-up or rejection of the submitted data) led to final approval of only 402 datasets (22.09%) (Fig. 2).

**Figure 2.**
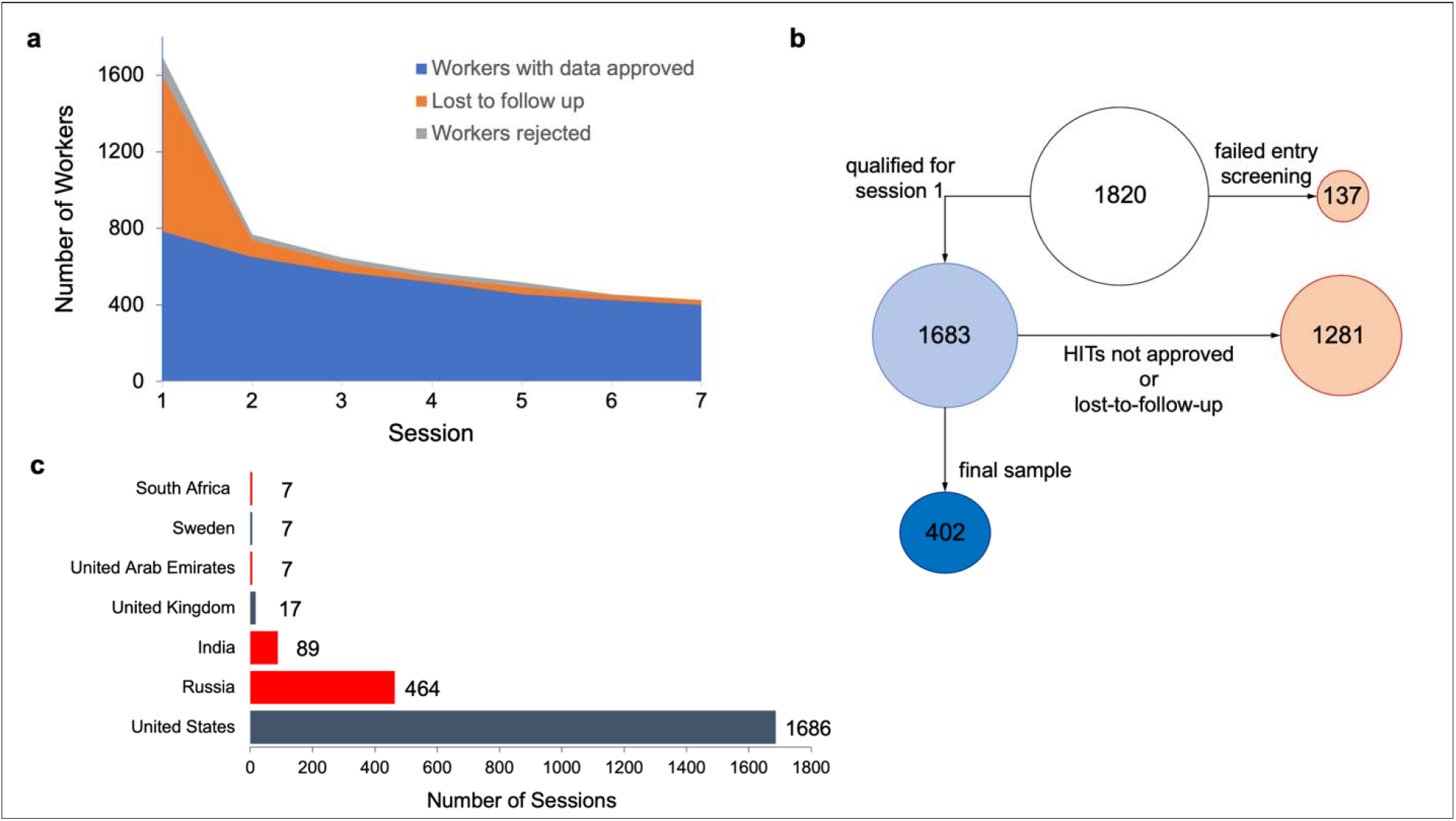
Sources of attrition and non-compliance over time. a) Worker attrition over 7 consecutive days. Colors reflect data loss due to rejection (gray, top), loss to follow up (orange, middle), and data retained (blue, bottom) over the seven-day experiment. **b) Flowchart of attrition from screening to final sample**. Diminishing circle sizes reflect loss of data from screening through session 1 and session 7. Circle colors reflect data loss due to non-compliance (orange, right), data retention (blue, left), and final sample (dark blue, bottom left). Numbers within circles reflect worker count. **c) Geolocation of workers over 2277 session**. Colors represent whether country from which data originated was on the list targeted for recruitment (dark blue) or not (red). Data and code to reproduce the figures 2a and c are available at https://doi.org/10.6084/m9.figshare.23499198.

To investigate a possible association between frame rate, lack of hardware acceleration, and geolocation of workers in the sample, we used an API call to www.ipify.org to record the participants’ internet protocol [IP] address during the experiment, and www.ipapi.com to geolocate the IP addresses during post-processing. Location information was available via IP address for 458 (25.16%) consented and screened participants across a total of 2277 sessions. We did not detect regional differences in hardware acceleration usage contributing to the study findings. Furthermore, the geolocation results indicate that workers submitted Mturk assignments from seven countries, specifically the United States, Russia, India, the United Kingdom, the United Arab Emirates, Sweden, and South Africa. While most submissions (74.04%) were from the United States, a large proportion of data (24.90%) originated from countries that were not on the approved list for the HIT on MTurk, namely Russia, India, United Arab Emirates, and South Africa (Fig. 2c, above).

In summary, in our longitudinal study, we identified two major sources of data fallibility. On the one hand, low and variable frame rates biased data quality, which could be mitigated through custom PsychoJS code but primarily affected sessions in which hardware acceleration was disabled. On the other hand, our longitudinal study was affected by high attrition, 77.91%, due to noncompliance and poor data quality leading to the rejection of submissions that exceeded the usual loss-to-follow-up. While geolocation data analysis did not reveal any clear patterns between frame rates and hardware acceleration utilization across regions, we received nearly a quarter of the submitted data from regions outside the locations explicitly specified as inclusion criteria on MTurk and for which we had obtained approval from the local Institutional Review Board.

## Discussion

Online crowdsourcing platforms such as MTurk provide significant advantages for experimental feasibility, for example, when aiming at larger than typical in-lab-collected sample sizes. Despite these benefits, the experiences gleaned through our study confirm known challenges and reveal significant additional ones that need to be carefully considered at all phases of online research. In this report, we identified two main sources of bias in time-sensitive data. The first involves technical shortcomings that affect data sampling, such as low and variable frame rates. The second pertains to human factors, particularly high attrition rates likely amplified by psychological factors such as motivation and competing attentional demands. These factors compromise the accuracy of response time measurements and can impact accurate inferences.

While our reflection is based on our experiences with MTurk, other online platforms (e.g., Prolific Academic [ProA], Crowdflower, Qualtrics, CloudResearch) are likely to face similar liabilities to varying degrees ^21–24^. All experimental work based on online crowdsourced data collection share extensive data *filtering* procedures and rejection of large amounts (>20%) of data ^8,10,25^. In the following, we discuss our findings in light of their consequences and propose potential solutions for the design of behavioral online experiments and the reporting of results, as well as technical developments required to ameliorate excessive burden on resources and wasting of collected data.

### Point 1. Technical Issues Contributing to Attrition: Hardware Acceleration and Frame Rate

Low and inconsistent frame rates, as demonstrated in our study, are major sources of inaccurate response time measurements, both within and across sessions. Taken together, the data show that: (a) frame rate tended to be lower and more variable with hardware acceleration off, (b) variability across sessions was high, regardless of hardware acceleration, and (c) within session, between-trial variability tended to be greater with hardware acceleration off. This compromise on the reliability and validity of study outcomes is primarily attributable to factors like participants’ software compatibility and hardware configurations, mainly hardware acceleration. Beyond hardware acceleration, other specific issues known to contribute to low and variable frame rates include the operating system used, with MacOS displaying less temporal precision than Windows or Ubuntu for visual stimuli, and the type of keyboard, as standard USB keyboards can introduce further latencies ^12^.

### Techniques to Manage Frame Rate Variability

To control these factors, some researchers suggest enforcing GPU hardware acceleration strictly in experiments where single-frame deviations or “dropped frames” are unacceptable ^14^. We argue that the effect of disabled hardware acceleration may be even more widespread, and not specifically limited to very high precision latency measurements, in the context of frame rate variability across longitudinal study designs. We implemented custom PsychoJS code as a solution to mitigate frame rate issues without GPU hardware acceleration by optimizing CPU rendering, although it’s unclear whether this software solution offers a comparable level of data reliability and consistency. Therefore, we concur with the recommendation to strictly enforce hardware acceleration not only when single-frame deviations are unacceptable but also when substantial variability in frame rate within and across sessions is also intolerable.

Other researchers recommend employing within-subject designs, limiting participants to a single browser, and restricting devices (e.g., operating system, laptop vs. desktop, USB vs. Bluetooth keyboard, etc.) used to reduce experimental noise ^11^. Contrary to this recommendation, our data suggest that frame rate variability within and across sessions remains a significant source of bias leading to unreliable data even in within-subject designs. Browser types do represent another significant limitation and cause of variability because it largely remains opaque how they, in general, and with each version update specifically, interact with hardware acceleration status or impact frame rate ^12,14^. One approach may be to restrict access to the experiment only through specified browsers, which again is another source of bias and limitation of participant diversity and does not account for unknown interactions between browser version updates and frame rate. Rather than strictly limiting participant hardware and software, implementing software solutions that control or standardize frame rates can help ensure consistent response time measurements within sessions, across sessions, and across participants.

Incorporating robust data screening and cleaning methods early on in a study can help mitigate the impact of frame rate variability detriment on study outcomes, for example, by defining acceptable frame rate thresholds for exclusion prior to beginning data collection. Considering that a standard computer monitor operates at a minimum of 30 fps, it is necessary for researchers to define a frame rate cutoff below this threshold; for example, 20 fps may strike a reasonable balance between hardware limitations and data reliability. Implementing a 20-fps frame rate, latency errors can range from 0 to 50 milliseconds resulting in a potential error of 50 milliseconds, which may be acceptable for some but not all experimental designs. It is important to note that the impact of frame rate on data quality may not be a simple binary issue but requires a more comprehensive exploration into whether there is a continuum of reliability as frame rate increases. Therefore, establishing clear guidelines for acceptable frame rates is essential to ensure the reliability of time-sensitive outcomes. However, researchers need to determine the acceptable margin of error based on their study design, decide whether hardware acceleration is a firm requirement, and then define a threshold of frame rate for exclusion, tailored to each individual study. While limiting study inclusion based on hardware acceleration and frame rate threshold a priori may limit sample diversity and bias results, ignoring these considerations may lead to excessive attrition and data wastage.

### Point 2. Human Factors Contributing to Attrition: Task Comprehension and Compliance

Our findings suggest that attrition rates in online crowdsourced studies can exceed 75% within a one-week timeframe—with the bulk of data loss from screening through session 1— substantially impacting study planning, budget, and time constraints. When screening is effective, however, much data loss can be prevented early on, leaving a remainder of participants committed to the longitudinal design. Besides the logistical implications of such high attrition, the resulting data loss clearly diminishes a study’s statistical power and increases the risk of biased results. Several factors likely contribute to this concern. First, the lack of face-to-face interaction might result in participants feeling less committed to completing the tasks. Second, typical research designs used to investigate procedural learning with high repetition rates clearly tax motivation and compound loss of participants’ interest over time. Third, the remuneration provided might not be perceived as adequate compensation for the time and effort required.

Another related challenge in navigating attrition is non-compliance in task execution. High HIT reject rates can occur due to participants not completing large portions of an experiment during a session or inappropriately performing a task (e.g., random key presses). Non-compliance such as this can largely be rooted out through screening and rejection during the first few sessions but may arise even in later sessions in workers whose performance has met criteria up until that point. In some cases, participants might be multitasking, putting additional load on their system, which leads to inconsistent response times that further contribute to frame rate variability. In these ways, technical and human factors interact in contributing to overall study attrition.

Notably, in the present study, most workers ignored multiple salient experimental prompts warning them to ensure that hardware acceleration was enabled and that all background processes were turned off prior to beginning each experimental session (at risk of submission rejection) via on-screen prompts and email reminders. Risk of rejection theoretically serves as a potent motivator for MTurk workers to comply with task requirements, although, as shown in our results, it was clearly insufficient in motivating the majority of workers to enable hardware acceleration prior to beginning each experimental session.

#### Strategies to Reduce Attrition due to Human Factors

One strategy to mitigate attrition and increase retention over several sessions is to use stepwise payment schedules as financial incentives. It is also important to ensure that study designs are engaging and not excessively laborious; this can also help maintain participant interest and commitment, which may, consequently, lead to the decision that a study design is not suited for online data collection.

Pre-screening participants for the likelihood of commitment based on past performance— as it is possible through limiting experimental access based on prior performance scores implemented on online platforms—and ensuring adequate hardware/software capacity are essential steps. This process helps identify the most likely participants to complete the study and provide high-quality data. While these measures may reduce attrition, they will obviously limit the diversity and extent of the worker pool accessed during recruitment.

Ensuring that participants understand the experimental requirements and excluding non-human entities, such as bots, are the two most essential goals of entry screening. Clear communication about study requirements is fundamental to enhancing participant motivation and reducing attrition due to dropout or misunderstanding of task instructions. Instructions should be concise and clear, with screening questions implemented to ensure comprehension. It is worth noting that stringent screening and exclusion based on inattentiveness to instructions can reduce sample sizes by over 50% across crowdsourcing platforms ^21^. Researchers may benefit from including attention checks during the screening that confirms understanding of hardware acceleration requirements and task comprehension. Alternatively, including a sample task would allow for the determination of hardware acceleration status and task comprehension prior to enrollment in the full study. Post-enrollment, implementing meticulous screening methods, such as daily data reviews based on transparent pre-defined quality criteria, is instrumental in assuring high data quality. As an example, performance metrics can be defined in advance for each experimental session, with stringency increasing as the study progresses.

In addition to these recommendations, we argue that researchers need to make an informed decision about online crowdsourcing platform usage, such as MTurk, and whether these companies provide reliable and trustworthy services to fulfill requirements for rigorous science. The structure and operations of online platforms directly influence the quality of data collected, and certain improvements are essential to enhance researchers’ trust in the data. In our view, the discussed problems we identified with MTurk recruitment underline the need for reform.

Firstly, there is the inability to ensure that workers are meeting geolocation requirements as advertised by MTurk. Despite these claims, our data show that although we specified thirty countries to recruit from through MTurk’s service, our geolocation analyses revealed that this filter was not effective in limiting recruitment to the defined regions. Claims regarding the efficacy of geolocation restriction are ineffectual given the ease and accessibility of Virtual Private Network [VPN] clients. At best, online platforms can enhance their control by not only restricting access to HITs based on geolocation during screening, but also for each session. While this would minimize variability, it could prevent access to workers who may be traveling between countries over the course of an experiment. The solutions in this regard are unclear. Another potential concern is bias in country representation due to the time of publishing, around noon EST in our case. Even though this timing should not impact which countries are recruited from, it likely influences the geographic distribution of participants and skews the representation of certain regions ^15^. Geolocation services need to be reliable, and this discrepancy presents a serious challenge for researchers who need to target specific geographic areas for their studies and may need to exclude data collected from outside the targeted regions.

Secondly, there’s the inability to confirm that workers only have one worker ID. We have come across a case where a worker admitted via email that they were enrolled in the study twice with different worker IDs. Although hopefully a rare occurrence, this compromises the validity of study outcomes and further erodes the trust researchers place in a platform. Other related issues concern the possible use of multiple accounts per worker, perhaps facilitated by using different IP addresses via a VPN, or multiple workers sharing the same worker ID. But, as Casey et al. (2017) noted, shared IP addresses do not necessarily indicate the same worker repeating a task. Researchers need to be aware of these limitations and implement their own software solutions to obtain detailed information about their participants, such as browser type and version, hardware specifications, and IP addresses. This information can help researchers identify potential issues that might affect the data quality and increase the burden of attrition and squandered resources.

### General Conclusions and Recommendations

We here report challenges and potential pitfalls intrinsic to online crowdsourced research and discuss measures to mitigate them. Monitoring frame rates and employing suitable software solutions to require hardware acceleration are critical to maintaining the accuracy of response time measurements. Strategies to promote task engagement and limit attrition are also essential, and could include sending email reminders, incorporating frequent attention and comprehension checks, leveraging the stigma of MTurk HIT rejection, and devising a thoughtful payment schedule that motivates participants with fair compensation while also considering the project’s budget and realistic attrition rates. This study underscores the need for researchers to be cognizant of the potential pitfalls associated with online crowdsourced research and to implement appropriate measures to ensure reliable and valid data collection. By acknowledging and addressing these challenges, researchers can continue to leverage the flexibility and appeal of online crowdsourcing platforms like MTurk, thereby facilitating effective psychophysics research. In conclusion, while there are challenges associated with conducting online crowdsourced research, there are also many potential solutions and areas for future research.

## Author Contributions

PAM, CF, and KFH designed the study, acquired and analyzed the data, wrote and edited the manuscript.

## Data availability statement

All source data to reproduce the figures are available under https://doi.org/10.6084/m9.figshare.23499195 and https://doi.org/10.6084/m9.figshare.23499198.

## Code availability statement

All code to reproduce the figures is provided alongside with the source data.

